# Bridging spatial scales in respiratory viral infection

**DOI:** 10.64898/2026.04.22.720070

**Authors:** Ulrik Hvid, Bjarke Frost Nielsen, Kim Sneppen

**Author notes:** Concept: UH, KS. Text: UH, BFN, KS. Background literature: UH, BFN, KS. Derivations: UH. Figures: UH. The authors declare no competing interests.

## Abstract

Respiratory viruses spread within the host through both local expansion and occasional long-range dissemination that seeds new infection foci. We present LEAP, an analytically tractable within-host model that captures this two-scale process by coupling local plaque growth to long-range seeding. The model reduces to an age-dependent branching process and yields a closed-form expression for the exponential growth rate during early infection. Using empirical data to parametrize the model, we find that productive dissemination requires only a small number of successful long-range seeding events per infected cell, with distinct values for SARS-CoV-2 and influenza A virus. LEAP further predicts that, in these well-adapted viruses, interferon-mediated restriction only weakly affects exponential growth, while remaining decisive for poorly adapted ones. More broadly, the model provides a flexible framework for experimentally testable predictions of early infection dynamics.

**Significance Statement:** Respiratory viral infection is an inherently spatial process, in which the virus must colonize large areas of the airways to optimize reproduction. Recent studies in animal models infected with influenza A or SARS-CoV-2 have documented long-range stochastic jumps of viral populations between distant regions of the respiratory tract. The emerging picture is one of two co-occurring spatial processes: slow, local plaque expansion and long-range seeding events that are rare but crucial to rapid colonization of the airways. We introduce LEAP (Lotka-Euler Airway Pathogen model), a simple mathematical within-host model that captures these two-scale dynamics by coupling local plaque growth to stochastic long-range seeding. Using LEAP, measurements from the petri dish can directly produce predictions of infection dynamics in the body.

---

During viral infection of the respiratory system, successful viral reproduction depends on the availability of cells that are 1) not already infected and 2) susceptible to infection. If a virion is released in an area of the lung with a high concentration of viruses and where, perhaps, antiviral signaling has begun to limit susceptibility, this incentivizes the virus to move to another part of the respiratory tract (RT). Fast dissemination will maximize viral shedding of the host, increasing the probability of infecting other individuals (1). Thus, viral colonization of the RT is a spatial process in which the virus must spread as widely as possible in the time window preceding activation of a systemic immune reaction involving widespread inflammation and recruitment of leukocytes (2). In the early stages of a successful respiratory infection, viral load grows exponentially (3–6). *In vitro* studies, however, in cell cultures can typically only show local neighbor-to-neighbor spreading of viruses (7), and such spatially constrained spread is incompatible with exponential growth. Although *in vitro* studies with liquid overlays can exhibit diffuse plaques and enhanced seeding of secondary plaques (7), this is still a far cry from the long-range transport seen *in vivo*. Indeed, recent studies (8–10) have made it clear that *in vivo* infection involves long-distance transport of viral particles, facilitated by air or bloodstream.

Along these lines, a recent study in PNAS clearly documented that infection with the H1N1 influenza A virus (IAV) in ferrets involves long-range stochastic jumps between regions of the respiratory system (8). Similarly, it has been shown that in mice coinfected with IAV strains that differ only by a fluorescent reporter protein, discrete islands of infection are established in the lungs (9, 10). Finally, SARS-CoV-2 establishes separate infection nodes throughout the RT of an infected hamster (11). Thus, the dynamics of viral proliferation involve an interplay between local spread between neighboring cells and long-range jumps of viral particles that can initiate new foci of infection (IF) (1). These observations provide clues to models that may explain, simultaneously, the spatially constrained growth observed *in vitro* and the observations from human challenge studies, which show exponential growth during the first 2-3 days of viral infection. For SARS-CoV-2, viral load has been shown to double every ≈ 2-3 hours in immune-naïve humans, peaking at 6.5 days post-infection (dpi) (3, 4). With IAV (H3N2) in humans, viral load can increase by 3 orders of magnitude between day 1 and 2 post-infection (12), corresponding to a similar doubling time to SARS-CoV-2, and peaking at 3 dpi.

Thus, we can think of *in vivo* infection as taking place on two separate spatial scales. The majority of published in-host dynamical infection models fall into two categories, each effectively describing one of these scales. Target-cell limited (TCL) models consist of Lotka-Volterra-style ordinary differential equations that treat the body as one well-mixed container, or a small number of organs that individually are well-mixed (13). Over the past three decades, such models have achieved considerable success in elucidating fundamental aspects of viral kinetics (14–19). In particular, landmark studies of HIV dynamics have shed light on key quantities such as the lifespans of productively and non-productively infected cells, the effects of (combination) antiretroviral therapy and the multiple phases of decline in plasma viremia associated with treatment (20). These achievements demonstrated how even highly idealized compartmental models – when combined with clinical and in vitro data – can yield important insights. Despite these successes, classical compartmental models relying on the assumption of homogeneous mixing often lack the granularity to discriminate between mechanisms. Furthermore, the acute diseases considered in this paper are fast-acting relative to HIV, likely making the well-mixed assumption less appropriate in this context. Other models have explicitly represented space, at the cost of being limited to describing local dissemination of viruses between neighboring cells (21–25). These models generally struggle to produce long periods of exponential growth, and are typically not analytically tractable.

In this study, we present a null-model of how the two-scale dynamics described above produce exponential growth, and how the growth rate depends on the two parameters that describe short- and long-range dynamics, respectively. We call it Lotka-Euler Airway Pathogen model or LEAP. Like TCL models, LEAP is analytically tractable, but allows for a more realistic representation of viral propagation, using a quasi-spatial representation. LEAP can take parameters directly from *in vitro* measurements, going beyond a phenomenological description that risks overfitting. We demonstrate a few simple extensions that increase biological realism while maintaining analytical clarity. With these extensions, we estimate the model parameter that is most difficult to measure directly, which we denote by *q*, the number of virions secreted by an infected cell that initiate new IF elsewhere, for both SARS-CoV-2 and IAV.

## Results

### The null-model

First we present the simplest version of LEAP. As we will argue later, this model has certain limitations in terms of biological realism, but it illustrates the central concepts of this study. We represent local spread as a circular viral plaque or focus of infection (IF) that grows at a constant radial speed *v*, as has been observed in *in vitro* studies (26– 28). These studies suggest a constant speed during the growth phase of the plaque on the order of 10 − 20 *µ*m*/*h ≈ 1 *d*_*c*_*/*h where *d*_*c*_ is the typical diameter of an epithelial tissue cell. Virions released within the locally growing plaque will then have a chance to be transported, via air or bloodstream, to another location in the RT. That new focus is assumed to be far enough away that the circles never overlap. The area *A* of an IF grows with its age *a* as

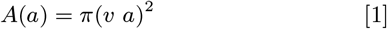

We use spatial units of *d*_*c*_ such that the area has units 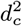 and simultaneously represents the number of infected cells. We use *q* to denote the number of new IF seeded per infected cell. Dimensionally, *q* is not a number per cell, but per unit area 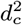. However, we assume a confluent monolayer with cell number density 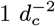. This is clearly a natural assumption in a monolayer culture, but justifiable even *in vivo*, since many common respiratory viruses predominantly infect and reproduce in the ciliated top layer of the epithelium (29–31), as seen in images of infected lung tissue (11).

The rate of new infections in an IF is the time-derivative of the area *A*, and the rate of production of new IF (birth rate *b*(*a*)) is

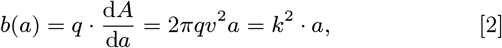

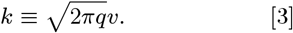

In other words, this system represents an age-dependent birth process in which the birth rate *b*(*a*) of each individual is proportional to its age. The analysis can then proceed without explicitly considering expanding circles, but instead taking individual IF reproducing at increasing rates as the building blocks. We call *b*(*a*) the kernel, and it contains many of the biological assumptions made in the model. In this study we will also analyze more realistic birth kernels (Fig. 1b).

**Fig. 1.**
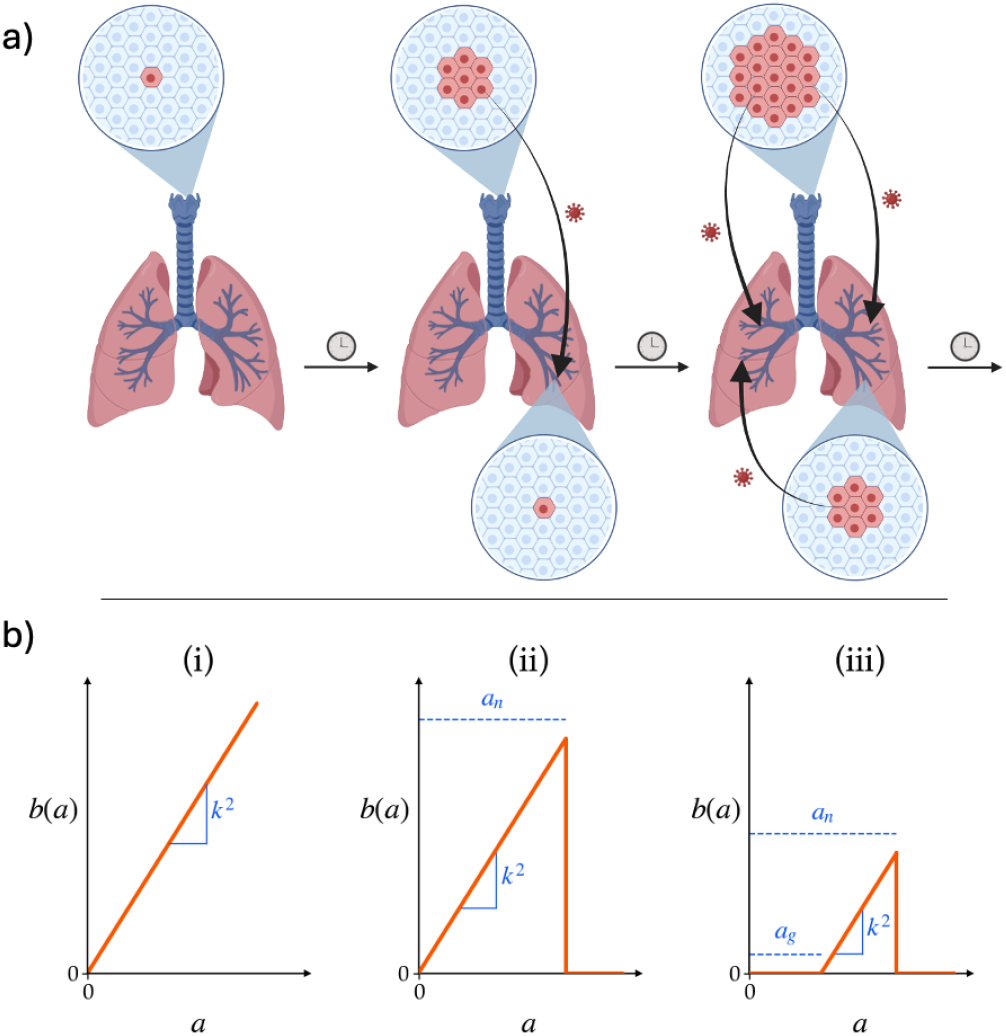
Model of the interplay of short- and long-range dynamics. **a)** The model consists of foci of infection that expand and replicate. Local expansion (red) happens at a constant radial speed. Each newly-infected cell seeds *q* (typically *<* 1) new foci of infection. Made with BioRender (32). **b)** The three different birth kernels used. (i) Simplest model. Linear growth starting at age *a* = 0 and growing without bounds. (ii) First expansion. Births go discontinuously to zero at *a* = *a*_*n*_. The actual input parameter is *p*_*n*_, while *a*_*n*_ is the age that satisfies *A*(*a*_*n*_) = 1*/p*_*n*_ (Eq. 12). (iii) In the final expansion we introduce a delay *a*_*g*_ between the seeding of a new focus and the time when it starts producing offspring. In all cases, 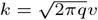.

From the kernel, we can derive the time evolution of the system as a whole. We will solve the simplest model (i) (Fig. 1b) in the main text and refer to Materials and Methods for the more involved derivations of models (ii) and (iii). Define *B*(*t*) as the total rate of births at time *t*. Then *B*(*t* − *a*)*b*(*a*) is the rate of births contributed at time *t* by individuals of age *a*. Since *B*(*t*) is the sum of all these contributions, plus the birthrate *b*(*t*) of the initial IF,

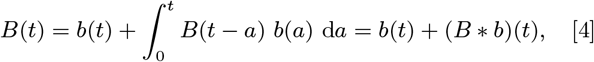

where ∗ is the Volterra convolution (33). We then take the Laplace transform, which converts convolutions to products (34):

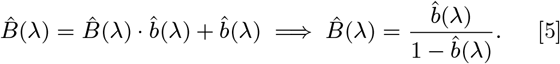

To recover *B*(*t*), we look for poles of the Laplace transform (35). In this case, a pole is a value *λ* that satisfies 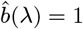, or

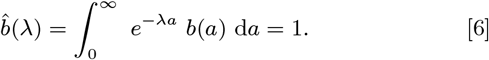

Eq. 6 is a version of the Lotka-Euler (LE) equation, which is commonly used to describe population growth when reproduction is age-dependent (36, 37). Using the kernel from Eq. 2, Eq. 6 becomes

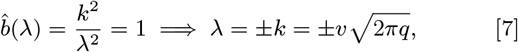

Since the positive growth rate dominates in the long run, we ignore the decaying *λ <* 0 solution. In conclusion,

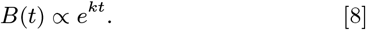

An exact derivation would yield *B*(*a*) = *λ* sinh(*λa*), but the procedure above – finding the largest *λ* solving the LE equation (the dominant pole of 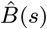 (35)) and assuming exponential growth – is easily generalizable to more complicated kernels and correctly identifies the growth rate that dominates after a transient. We assume 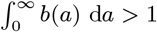 in all cases, since this is required for a growing population and ensures the existence of a positive *λ*.

After the transient, the population of IF settles into an exponential age distribution

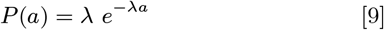

yielding the mean values of age and area, respectively,

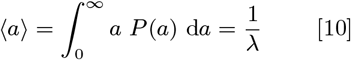

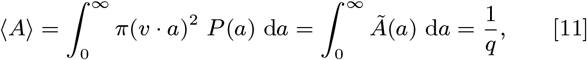

using Eq. 7. Here, *Ã*(*a*) ≡ *A*(*a*) · *P* (*a*) is the contribution to the mean area due to plaques of age *a*, and it is plotted in Fig. 2a. Note that none of these results require knowledge of the viral generation time, the burst size of an infected cell, or the diffusive properties of viral particles in epithelial tissue. Instead, these properties combine into the emergent expansion speed *v*, which can be estimated from *in vitro* measurements.

**Fig. 2.**
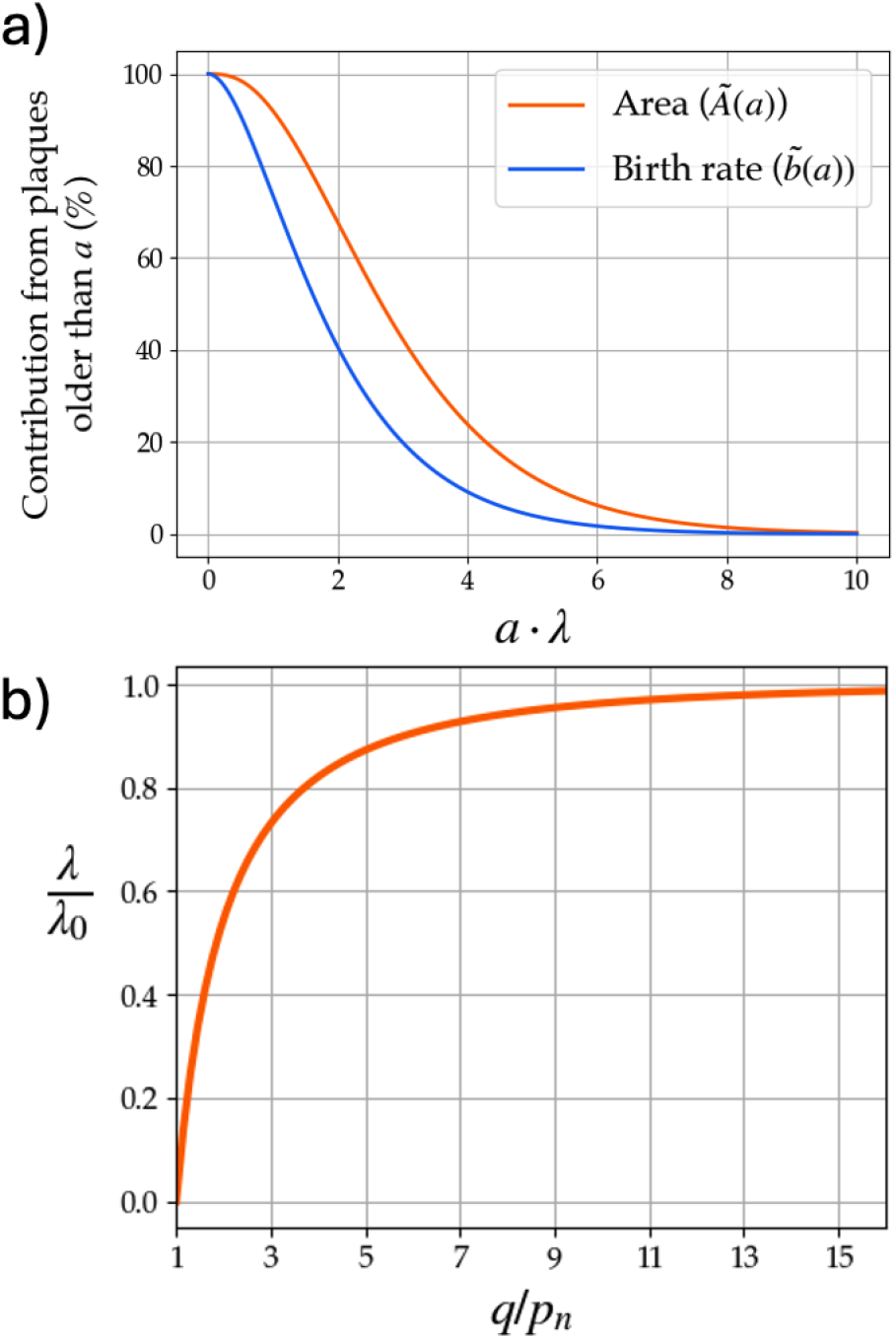
Diminishing returns from cytokine signaling in limiting viral growth rate. **a)** Survival function of *Ã* (orange) and 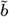 (blue). *Ã*(*a*) ≡ *P* (*a*) · *A*(*a*) and *b*(*a*) ≡ *P* (*a*) · *b*(*a*) are the relative contribution by IF of age *a* to the total infected area and total birth rate, respectively. We see that IF with *a > /λ* contribute less than 10% of total infected area and ≈ 1% of new births. **b)** Effect of IFN signaling on growth rate, measured as the actual growth rate *λ* divided by the counterfactual growth rate *λ*_0_ in the absence of IFN. The asymptotic shape shows that, if *p*_*n*_ ≪ *q*, the virus cannot increase growth rate further by decreasing *p*_*n*_.

### Introducing IFN response

The LE equation typically includes a death factor (37, 38) that brings the birth rate down to zero with increasing age, but we have so far allowed *b*(*a*) to grow unbounded. As such, the reproduction number of a plaque is infinite in the null-model. Intuitively, one might guess that this implies super-exponential growth, but the analysis above shows that this is not the case. The reason is that the significance of older IF gets “diluted” among the much greater number of younger offspring (Fig. 2a).

However, real IF do not grow infinitely, and we should examine how problematic this assumption is. The first extension of the model ((ii), Fig. 1b) has IF stop their growth due to the first line of immune defense, namely cytokine signaling and the resulting antiviral state in infected tissue cells. It is well documented that interferon (IFN)-competence in host cells can have the effect of halting expansion of viral plaques (24, 39), and in differentiated air-liquid-interface (ALI) cultures of primary human bronchial epithelial cells infected with SARS-CoV-2, spatial expansion stops after ⪅ 3 days (40). Just a single IFN-producing cell can likely activate antiviral defenses in a large radius around the infected area (27, 41).

We denote the probability that an infected cell secretes IFN and stops expansion of the local IF by *p*_*n*_. An order-of-magnitude estimate of *p*_*n*_ for SARS-CoV-2 can be inferred from experiments in (40), since in those cultures infection typically stops after ≈ 10^4^ infected cells, suggesting *p*_*n*_ ≈ 10^−4^. For IAV, which is far less competent at antagonizing antiviral responses (42), *p*_*n*_ has been estimated 50 times higher, 5 · 10^−3^ (43). This may be an upper bound as it does not correct for the paracrine feedback loop in which IFN stimulation increases the probability of IFN secretion (44).

Model (ii) is deterministic and assumes that IF expansion always stops when the area is exactly 1*/p*_*n*_. Stochasticity in IFN activation can be represented more accurately by including an exponential distribution in the integral, but the resulting equation is far more complicated (see supplementary material). According to Eq. 1, *A*(*a*) = 1*/p*_*n*_ when

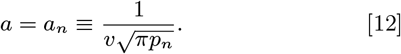

Now, *b*(*a*) = *k*^2^ · *a* for *a < a*_*n*_ and *b*(*a*) = 0 for *a > a*_*n*_. The growth rate is found by solving the LE equation (Eq. 6) but taking the integral only up to *a*_*n*_. The solution is the transcendental equation

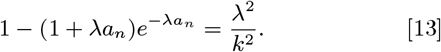

This equation shows the dependence of the growth rate on *a*_*n*_ and thus *p*_*n*_. When *a*_*n*_ → ∞, we recover Eq. 7. To evaluate the effect of IFN signaling on growth rate, we define

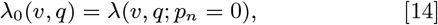

i.e. the growth rate that would be observed in an equivalent system with no IFN response. Then we can quantify the effect of IFN response by *λ/λ*_0_, as in Fig. 2b. We find a steeply rising *λ/λ*_0_ for *q/p*_*n*_ ≈ 1 beyond which it asymptotes to 1. Evidently, decreasing *p*_*n*_ by antagonizing antiviral signaling yields diminishing returns for the virus. In general, if *p*_*n*_ ⪅ *q/*10, a further reduction in *p*_*n*_ hardly affects the growth rate (Fig. 2b). The reason is apparent from Fig. 2a; although the oldest IF are, individually, the largest and most productive, the exponential age distribution (Eq. 9) implies that they are much less abundant than the younger IF. As a result, the total infected area and the birth rate are dominated by younger, smaller plaques with age *a <* 10*/λ*. This means that unless IFN kicks in at *a*_*n*_ ≪ 10*/λ*, the effect on growth dynamics will be negligible.

### The full model - time delay

In terms of biological realism, the main drawback of the models presented so far is the fact that a newly seeded IF starts producing offspring immediately after birth. This is problematic because the time it takes for the seed cell to even start transcribing viral genes on a large scale is greater than the eventual doubling time of the system, e.g., ≈ 6 h for SARS-CoV-2 (45) vs. 2-3 h of doubling time (3, 4). Knowing this, the result from Fig. 2a that plaques with *a <* 1.5*/λ* ≈ 6 h account for 50% of the birth rate cannot be remotely correct. Therefore, we introduce a delay *a*_*g*_ in the growth of each IF, leading to the birth kernel (iii) shown in Fig. 1b. The mathematics of this full model is derived in Materials and Methods, where we show that the dynamics are fully contained in the equations

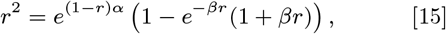

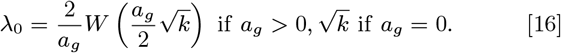

with *r* ≡ *λ/λ*_0_, *α* ≡ *a*_*g*_*λ*_0_, *β* ≡ (*a*_*n*_ − *a*_*g*_)*λ*_0_ and *W* (·) the Lambert W function (46). Fig. 3 shows how the dynamics depend on *a*_*g*_ . If we fix *q* and allow *λ*_0_ to change according to Eq. 16, we get Fig. 3a. We see that *a*_*g*_ decreases the growth rate, which has the effect of widening the *Ã*-distribution, increasing the contribution of older and larger IF. Interestingly, even in the full model, ⟨*A*⟩ = ∫ *Ã* d*a* = *q*^−1^, so the area under the curve remains constant (Fig. 3b, proven in supplementary material).

**Fig. 3.**
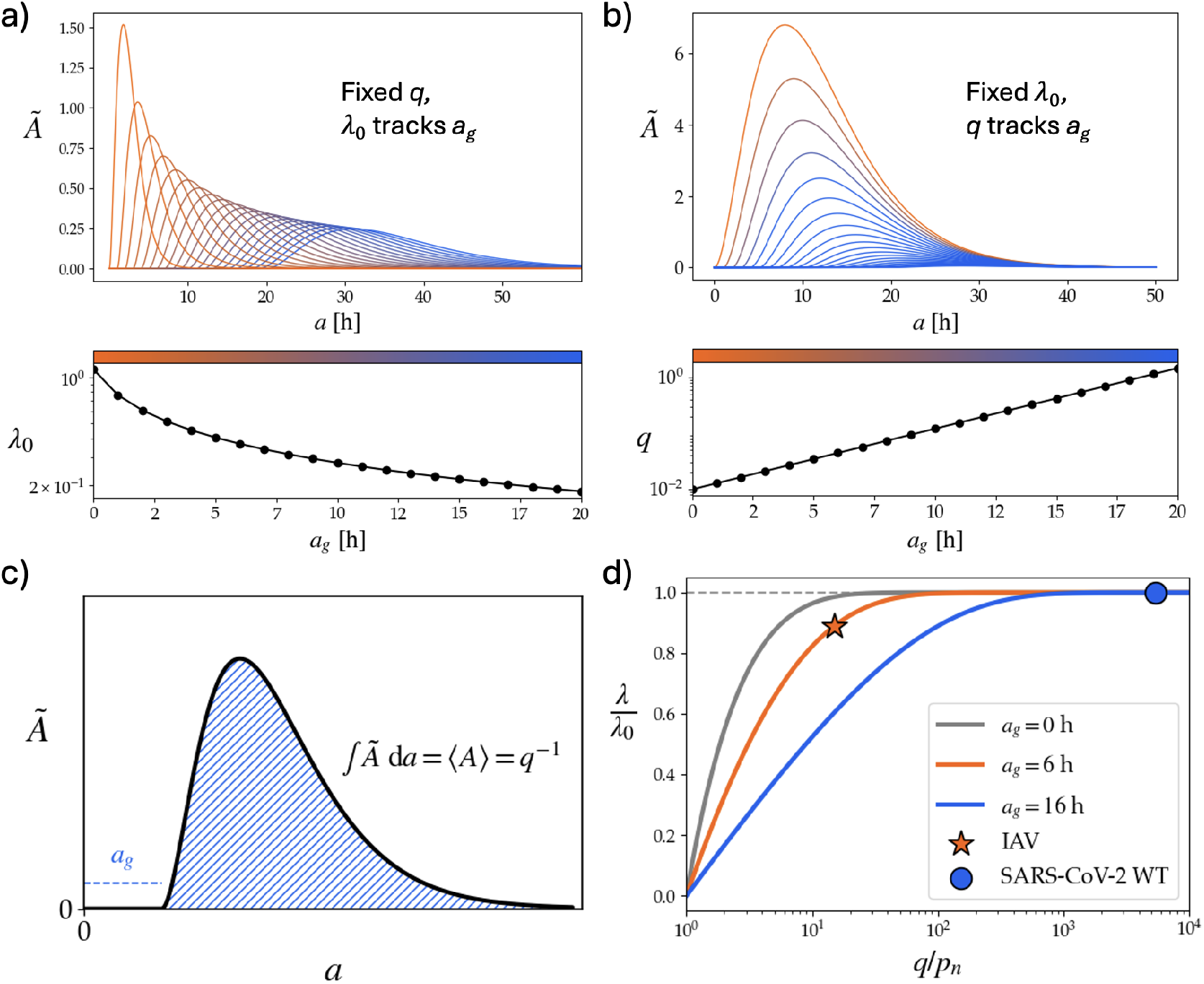
Introducing time delay in IF growth alters dynamics. **a)** Distributions of the age-dependent area contribution *Ã*(*a*) for various values of the delay *a*_*g*_ and fixed propagation speed *v* = 1 and jump parameter *q* = 0.2. Increasing *a*_*g*_ decreases *λ*_0_ according to Eq. 16. **b)** When keeping *λ*_0_ fixed by adjusting *q*, the area under the curve shrinks as *q* increases according to Eq. 28. Note that *q* must increase exponentially with *a*_*g*_ to ensure a constant *λ*_0_ . Here, *v* = 1, *λ*_0_ = 0.25. **c)** Schematic of the relation between the *Ã* distribution, the mean area and the jump parameter. As shown in the supplementary material, the area under the curve is equal to *q*^−1^. **d)** Effects of IFN on the exponential growth rate. The effect is quantified by *λ/λ*_0_, the ratio of the actual growth rate to the growth rate that would be observed in the absence of IFN signaling.

Instead of keeping *q* fixed, we can also let it track *a*_*g*_ to keep *λ*_0_ constant. The result is shown in Fig. 3b, where *λ*_0_ = 0.25 h^−1^. The bottom panel shows that this requires *q* to increase exponentially with *a*_*g*_ (Eq. 28). The increasing value of *q* is expressed as a decreasing area under the *Ã*-curve as *a*_*g*_ increases, according to Eq. 28. The result of diminishing returns, illustrated in Fig. 2b, remains valid, though the range over which the IFN has a measurable influence increases dramatically, Fig. 3d.

### Estimating the jump parameter

To estimate *q*, we need to fix all other parameters, as well as the resulting growth-rate *λ*. The parameters for SARS-CoV-2 and IAV are shown in Table 1. To determine *a*_*g*_, we looked at one-step growth curves, i.e., viral titers from cultures infected at high multiplicity of infection, to represent the output of just one viral generation. We set *a*_*g*_ equal to the time when the viral output of the seed-cell has the biggest impact on overall growth. For SARS-CoV-2 the curve has a complex shape, with a small output already at ≈ 6 hpi, but increasing by many orders of magnitude over the next 36 h. (47). To determine the time at which the output of the seed cell is most significant for growth, we discount the late offspring with the factor *e*^−*λt*^ – in line with the logic of the LE equation – and locate the peak, (supplementary fig. S1). For the wild-type (WT) variant, there is a clear peak at 16 hpi, and for Delta there is a peak at 12 hpi. Since the exponential growth rate was measured for WT SARS-CoV-2 (3, 4), we use 16 hpi. For the IAV the curve is simpler, as the shedding rate increases almost stepwise at around 6 hpi (48), so we set *a*_*g*_ = 6 h for IAV.

**Table 1.**
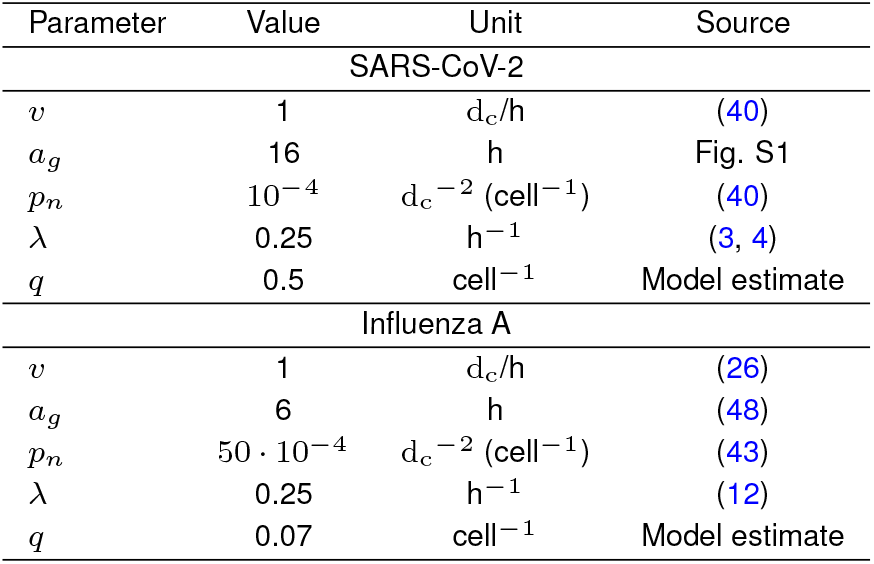
Model parameters for SARS-CoV-2 and IAV.

| Parameter | Value | Unit | Source |
| --- | --- | --- | --- |
| **SARS-CoV-2** |  |  |  |
| $v$ | 1 | $d_c/h$ | (40) |
| $a_g$ | 16 | h | Fig. S1 |
| $p_n$ | $10^{-4}$ | $d_c^{-2} (\text{cell}^{-1})$ | (40) |
| $\lambda$ | 0.25 | $\text{h}^{-1}$ | (3, 4) |
| $q$ | 0.5 | $\text{cell}^{-1}$ | Model estimate |
| **Influenza A** |  |  |  |
| $v$ | 1 | $d_c/h$ | (26) |
| $a_g$ | 6 | h | (48) |
| $p_n$ | $50 \cdot 10^{-4}$ | $d_c^{-2} (\text{cell}^{-1})$ | (43) |
| $\lambda$ | 0.25 | $\text{h}^{-1}$ | (12) |
| $q$ | 0.07 | $\text{cell}^{-1}$ | Model estimate |

**Table 2.**
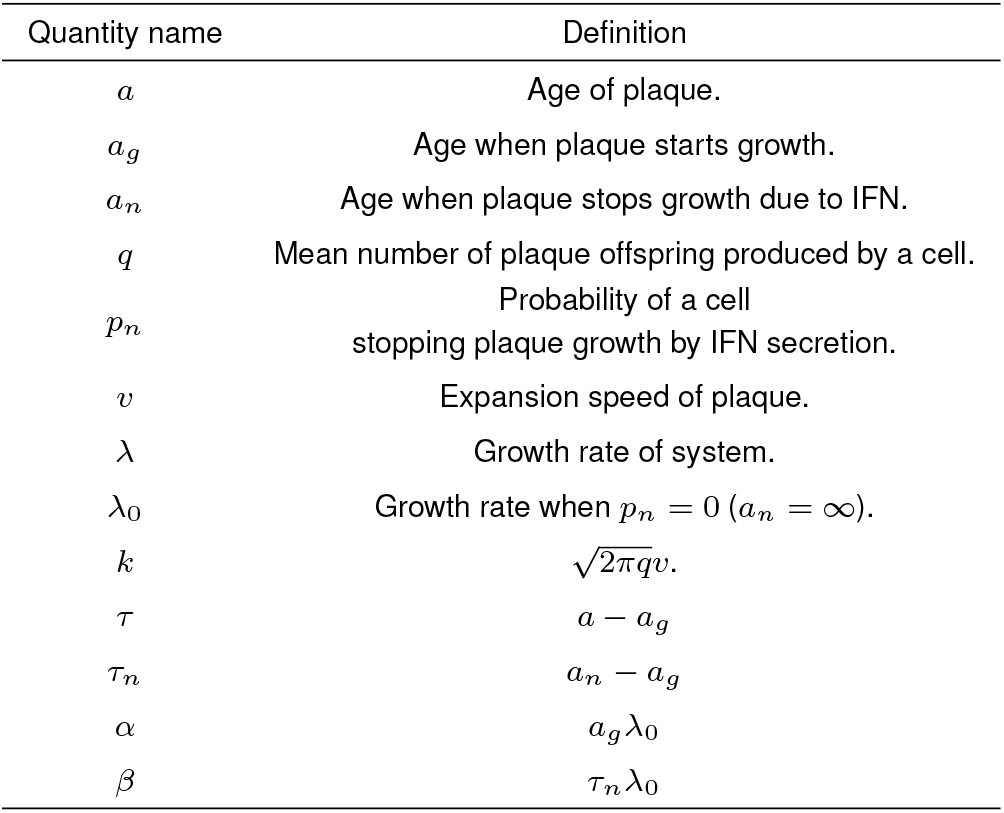
Definitions of quantities used.

To determine *q*, we solve Eq. 27. We get 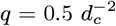 for SARS-CoV-2 WT. As a sanity check, the time *a*_*n*_ it takes for a plaque to reach *A*(*a*_*n*_) = 1*/p*_*n*_ and stop growing is 72 h in agreement with the ≈ 3 days it takes for an IF to stop growing in ALI cultures (40). For IAV, we get *q* = 0.07. The fact that the jump rate is much lower for IAV is not surprising, since it is well known that individual IAV particles are typically not able to complete infection alone (49). One of the reasons is that the majority have deleterious mutations (43). In addition, these mutations are strongly correlated with increased activation of antiviral responses, helping to explain the higher *p*_*n*_ for IAV. Conceptually, the jump parameter can be decomposed into several independent factors:

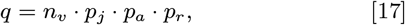

where *n*_*v*_ is the number of fully infectious progeny per cell, *p*_*j*_ is the probability that a virion is transported to a distant location in the RT and penetrates the mucous membrane there, *p*_*a*_ is the probability that such a virion adsorbs into a cell at that location, and *p*_*r*_ is the probability of completing reproductive infection there. Then, the high fraction of defective IAV particles (70-99% (49)) would reduce *q* via *p*_*r*_ .

Interestingly, we predict that for IAV, *p*_*n*_ is in the regime of increasing returns, as opposed to SARS-CoV-2 (Fig. 3d). If *p*_*n*_ were doubled, this would reduce *λ* from 0.25 h^−1^ to 0.21 h^−1^, which compounds into a difference of a factor of ≈ 7 over 48 h.

## Discussion

This paper introduces LEAP, an analytically tractable model that couples local plaque expansion with stochastic long-range seeding events. The two-scale formulation reduces to an age-dependent branching process and predicts an exponential growth rate that is remarkably insensitive to the fate of larger plaques. Two extensions of the model – a cap on IF size via the parameter *p*_*n*_ and an IF growth delay *a*_*g*_ (Fig. 1b) – allowed for a more detailed biological analysis and yielded an estimate for the number *q* of jumping virions per cell on the order of 0.5 for SARS-CoV-2 and 0.07 for IAV. The model is useful in its simplicity, but should be applied with due respect for limitations. The estimate of *q* was highly dependent on the estimated growth delay *a*_*g*_ (Fig. 3b).

In traditional TCL models, the variables represented describe total scalar amounts of virus, infected/susceptible cells, IFN, etc. in the body. A major benefit of the LEAP model is that it also predicts the distribution of these quantities between different infection sites, and thus enables additional falsifiable predictions. For example, we predict that IFN deficiency will not affect the exponential growth rate for SARS-CoV-2, and will only modestly affect that of IAV (Fig. 3d). Previous studies of SARS-CoV-2 in mouse models give contradictory reports, with one study (50) showing a minimal effect of IFN deficiency on viral load at 2 dpi and another (51) showing a substantial difference. A more focused test of the model would require reparameterizing it to mice and making sure to only compare titers within the exponential phase, which may be shorter for mice than for humans, given their smaller lungs.

The subdivision of *q* into different factors (Eq. 17), also points to falsifiable predictions. Both *n*_*v*_ (burst size), *p*_*a*_ (probability of adsorption) and *p*_*r*_ (probability of reproductive infection) are measurable quantities that can be tuned with genetic modification, and our model makes concrete predictions as to how such changes should affect the growth rate *in vivo*.

At first glance, the result that the growth rate of SARS-CoV-2 is insensitive to the fraction of IFN secreting cells *p*_*n*_ seems at odds with the well-established notion that the ability of the virus to interfere with immune signaling of host cells is the key to its success. SARS-CoV-2, in particular, is known to benefit from the ability to substantially delay IFN induction (42, 52, 53). However, there may be no contradiction here, as the molecular pathways leading to IFN secretion are simultaneously associated with the secretion of cytokines that activate the leukocytic immune system (54). The mouse experiments in (50) showed precisely that, while viral load was unaffected by knocking out the receptor that binds type-I IFN receptors, that mutation did significantly affect recruitment of inflammatory immune cells. Thus, while the virus’ returns on antagonizing IFN secretion may be diminishing in terms of increasing its growth rate, it may still be able to extend its growth phase by offsetting the activation of the next immune layer. Finally, the conclusion of diminishing returns from antagonizing IFN relates only to well-adapted viruses that already do this effectively. Any respiratory virus is subject to the condition that, to grow, it must have *p*_*n*_ *< q*, otherwise a plaque will be halted before it can reproduce. It is plausible that IFN secretion from structural tissue continually prevents growth of viruses that are not adapted to humans but still enter our body. For instance, Newcastle Disease Virus (NDV) – a bird virus – triggers IFN secretion with as much as 25% probability (55), which probably renders successful infection of humans impossible. Indeed, knocking out IFN receptors in mouse macrophages drastically increases viral output of NDV (56).

The simplicity of LEAP relies on the fact that it does not represent gradations of jump distances, meaning that it can model biphasic spatial spread without introducing explicit spatial coordinates. This is the core assumption of the model; other assumptions can be relaxed by changing the shape of the birth kernel and, if necessary, solving the LE equation numerically. Indeed, the “space-free” assumption is extremely useful but has its limitations, as it may be that most jumps are of intermediate distance. The spatial correlations observed in (8) show that the dynamics are not entirely space-free. Nonetheless, that study does also document very long jumps, and due to the branched structure of the lung, it is possible for two plaques to be a short Euclidean distance from each other but still widely separated in terms of distance along the lung surface. In the future, we will work on explicitly representing viral proliferation in the branching geometry of the lung.

Regardless of the distribution of jump distances, the assumption of no overlap is increasingly violated as the amount of susceptible tissue decreases with infection. In a hamster infected with fluorescent SARS-CoV-2, the lungs appear nearly saturated already at 2 dpi (11). This raises the question of whether exponential growth is first curbed by immune responses or by lack of uninfected lung regions. In any case, the LEAP model claims only to describe the exponential growth phase.

Another simplification was to assume that IF expand isotropically with constant radial speed. In the lungs, the activity of ciliated cells causes a directed upward flow of the mucus layer, and a viral IF is likely to spread anisotropically, along elongated structures. However, an ALI study (40) of this movement indicates a characteristic speed similar to the one used in this analysis, and the precise geometry (expanding circle vs. expanding line) is not crucial for our analysis. *In vitro* studies also show that the geometry of IF is affected by the viscosity of the overlay (analogously, the mucus layer) (7, 26–28).

For virus types that shed their offspring from the live host cell over a period, e.g., influenza, SARS-CoV-2 or Vesicular stomatitis virus (57–59), the IF expansion speed *v* is highly sensitive to the first viruses that leave an infected cell (60). For SARS-CoV-2, for example, infected cells can release some progeny virus as early as 6 hpi, but release the vast majority much later (supplementary fig. S1, (47)). Possibly, this provides the strategic advantage of a temporal sub-division, with early release increasing *v* while bulk release increases *q*. The relatively slow expansion speed (4 *µ*m/h) measured for adenovirus (61) may speak to the disadvantage of a bursting the host cell. To inform future iterations of this model, it would be useful to experimentally test the effect on *v* of mutations that affect viral reproduction.

We showed that, in order to keep *λ*_0_ constant, the jump probability *q* has to increase exponentially relative to the delay *a*_*g*_ . While *q* is largely determined by the environment (the tissue in which the virus spreads), viruses may also evolve to increase their *q* value. For example, IAV has been shown to display receptor-binding hemagglutinin (HA) and receptor-cleaving neuraminidase (NA) proteins in a polarized fashion, facilitating their movement through sialic acid-rich host mucus (62, 63). In terms of the decomposition in Eq. 17, this asymmetric protein organization increases *p*_*j*_ . Similarly, different viruses may respond differently to mucociliary clearance and even benefit from it at times (40).

Future studies might extend our two-tier model to bridge the gap to subsequent layers of the immune system, beginning with the recruitment of NK cells and other leukocytes. Such an extension raises questions about the spatial precision of the next layer of the immune system. Perhaps each individual plaque faces some probability of being discovered by leukocytes that may start a signaling cascade, stopping the infection in its tracks. Alternatively, the next layer may be best modeled as purely systemic, being activated only once the *total* amount of IFN in the RT reaches some threshold. These two sets of assumptions produce entirely different models and predictions. The observation from a SARS-CoV-2 study (2) that immune cell activation near the inoculation site within 1 dpi is associated with mild disease seems to support the former assumption. In the LEAP model, probability of an IF activating immune cells locally could be wrapped in the sub-parameter *p*_*r*_ or incorporated as a death factor in the kernel.

In conclusion, this study offers a new framework for thinking about viral spread in the host. The model presented is parsimonious, yet mechanistically detailed and, we believe, empirically well-founded. Via the birth kernel, the model can be flexibly adjusted to different pathogens or future experimental findings. Given the observation that the upper airways are perpetually teeming with viruses that fail to establish productive infection (64), a detailed understanding of the early stages of infection is a necessary complement to the ample literature on the mechanics of humoral immunity. We believe that the growing evidence that viral spread in the RT is intrinsically spatial calls for new modeling frameworks. The model presented here is an attempt at such a framework and may open new avenues for virology.

## Materials and Methods

### Introducing time delay

In this section we derive the most detailed version of the model, including both delay in growth *a*_*g*_ and a maximum age *a*_*n*_ at which plaques stop growing. Assume a delay age *a*_*g*_ . The probability per cell of being an IFN responder is *p*_*n*_, and IFN is released at age *a*_*n*_, solving

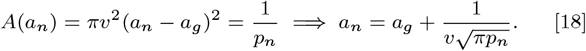

The typical age at which a plaque spawns a new focus of infection can be found similarly,

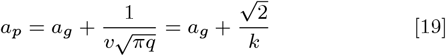

with *k* still defined as Eq. 3. Without IFN, the birth-kernel is

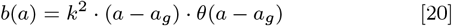

The Heaviside function is equivalent to introducing a lower bound in the integral of the Lotka-Euler equation. Thus, the growth rate in the absence of IFN, *λ*_0_, solves

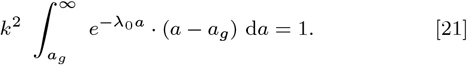

This can be simplified by substituting *a* for *τ* ≡ *a* − *a*_*g*_, so we get

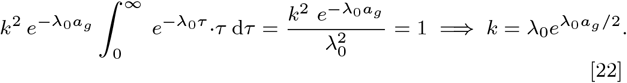

Eq. 22 is a transcendental equation for *λ*_0_ which can be written in terms of the Lambert W function (46),

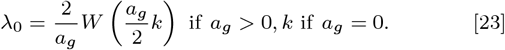

Including IFN-activity at *a* = *a*_*n*_, the Lotka-Euler equation gets an upper bound *τ*_*n*_ ≡ *a*_*n*_ − *a*_*g*_ :

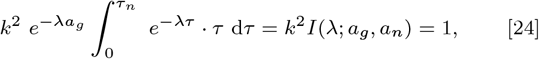

with

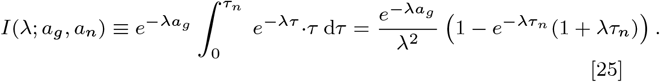

Now, using 22, the Lotka-Euler equation becomes

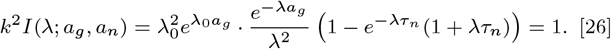

Notice that the jump probability *q*, which is the most difficult parameter to observe, is contained in *k* (Eq. 3) but not in the integral and can thus be estimated by

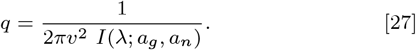

In the IFN-free case, the relation between *q* and *λ* = *λ*_0_ becomes quite simple:

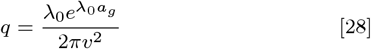

Eq. 26 is quite messy, with many interdependent variables, so we next try to reduce it to the minimum number of independent variables. Defining 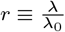, it can be written as

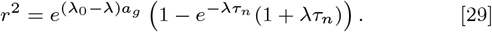

Factoring out *λ*_0_,

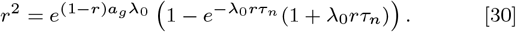

Finally, defining two dimensionless quantities *α* ≡ *λ*_0_*a*_*g*_ and *β* ≡ *λ*_0_*τ*_*n*_ this condenses to

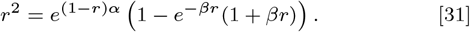

Thus, the relative effect *r* of introducing interferon depends on two independent variables *α* and *β*. The first is the delay before plaques start growing, the second is the time window in which the plaque is allowed to grow, both measured in terms of the characteristic time 1*/λ*_0_. Note that Eq. 31 is transcendental in *λ* and must be solved numerically.

We would like to be able to model the dynamics in terms of the more concrete biological parameter *p*_*n*_, rather than *a*_*n*_, which is more system-specific. To do this, define *τ*_*p*_ ≡ *a*_*p*_ − *a*_*g*_, similarly to *τ*_*n*_. Then

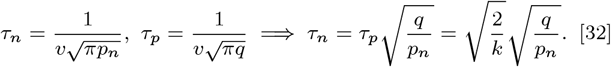

But since 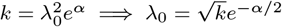 (Eq. 22),

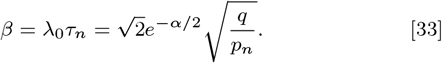

Thus, finally, we have shown that the relative significance *r* of IFN-signaling is uniquely determined by a the combination of *α* (time delay in units of the reproduction time-scale) and 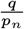, the ratio of jump parameter to the IFN first-response rate. For this reason, the curves in Fig. 3d are general and not reliant on specific absolute parameter values.

ChatGPT 5 and Claude Sonnet 4 aided in mathematical derivations and literature search. All mathematics in the article has been worked through by the authors and all cited sources have been read by the authors.

## Supporting information

Supplemental figures and derivations

## Funding

This research was funded by the Danish National Research Foundation (Grant no. DNRF170). BFN acknowledges financial support from the Carlsberg Foundation (Grant no. CF24-1337).

## Data and materials availability

Simulation code and data is available at the public Github repository (65).

## Notes

https://github.com/UlrikHvid/LEAP---Lotka-Euler-Airway-Pathogen-model

### Competing Interest Statement

The authors have declared no competing interest.

### Summary of Updates

The manuscript has been revised with a more descriptive title, a clearer abstract and a significance statement.

## References

1. ME Gallagher, CB Brooke, R Ke, K Koelle, Causes and consequences of spatial within-host viral spread. Viruses 10, 627 (2018).

2. RG Lindeboom, et al., Human sars-cov-2 challenge uncovers local and systemic response dynamics. Nature 631, 189–198 (2024).

3. SA Iyaniwura, et al., The kinetics of sars-cov-2 infection based on a human challenge study. Proc. Natl. Acad. Sci. 121 (2024).

4. B Killingley, et al., Safety, tolerability and viral kinetics during sars-cov-2 human challenge in young adults. Nat. Medicine 28, 1031–1041 (2022).

5. TG Evans, et al., Assessment of cd8+ t-cell mediated immunity in an influenza a(h3n2) human challenge model in belgium: a single centre, randomised, double-blind phase 2 study. The Lancet Microbe 5, 645–654 (2024).

6. H Miao, et al., Quantifying the early immune response and adaptive immune response kinetics in mice infected with influenza a virus. J. Virol. 84, 6687–6698 (2010).

7. Y Zhu, J Yin, A quantitative comet assay: imaging and analysis of virus plaques formed with a liquid overlay. J. virological methods 139, 100–102 (2007).

8. LM Ferreri, et al., Dispersal of influenza virus populations within the respiratory tract shapes their evolutionary potential. Proc. Natl. Acad. Sci. 122 (2025).

9. A Sims, et al., Superinfection exclusion creates spatially distinct influenza virus populations. PLOS Biol. 21, e3001941 (2023).

10. S Fukuyama, et al., Multi-spectral fluorescent reporter influenza viruses (color-flu) as powerful tools for in vivo studies. Nat. Commun. 6 (2015).

11. Bagato, et al., Spatiotemporal analysis of sars-cov-2 infection reveals an expansive wave of monocyte-derived macrophages associated with vascular damage and virus clearance in hamster lungs. Microbiol. Spectr. 12 (2024).

12. N Shetty, et al., Influenza virus infection and aerosol shedding kinetics in a controlled human infection model. J. Virol. 98 (2024).

13. D Higgins, et al., Introducing a framework for within-host dynamics and mutations modelling of h5n1 influenza infection in humans. J. The Royal Soc. Interface 22 (2025).

14. S Bonhoeffer, RM May, GM Shaw, MA Nowak, Virus dynamics and drug therapy. Proc. Natl. Acad. Sci. 94, 6971–6976 (1997).

15. AS Perelson, AU Neumann, M Markowitz, JM Leonard, DD Ho, Hiv-1 dynamics in vivo: Virion clearance rate, infected cell life-span, and viral generation time. Science 271, 1582–1586 (1996).

16. P Baccam, C Beauchemin, CA Macken, FG Hayden, AS Perelson, Kinetics of influenza a virus infection in humans. J. virology 80, 7590–7599 (2006).

17. A Handel, IM Longini, R Antia, Towards a quantitative understanding of the within-host dynamics of influenza a infections. J. Royal Soc. Interface 7, 35–47 (2010).

18. CA Beauchemin, A Handel, A review of mathematical models of influenza a infections within a host or cell culture: lessons learned and challenges ahead. BMC public health 11, S7 (2011).

19. SM Ciupe, JM Heffernan, In-host modeling. Infect. Dis. Model. 2, 188–202 (2017).

20. AS Perelson, RM Ribeiro, Modeling the within-host dynamics of hiv infection. BMC biology 11, 96 (2013).

21. C Quirouette, NP Younis, MB Reddy, CAA Beauchemin, A mathematical model describing the localization and spread of influenza a virus infection within the human respiratory tract. PLOS Comput. Biol. 16, e1007705 (2020).

22. C Beauchemin, J Samuel, J Tuszynski, A simple cellular automaton model for influenza a viral infections. J. Theor. Biol. 232, 223–234 (2005).

23. GA Funk, VA Jansen, S Bonhoeffer, T Killingback, Spatial models of virus-immune dynamics. J. Theor. Biol. 233, 221–236 (2005).

24. TJ Howat, C Barreca, P O’Hare, JR Gog, BT Grenfell, Modelling dynamics of the type i interferon response toin vitroviral infection. J. The Royal Soc. Interface 3, 699–709 (2006).

25. X Xu, BF Nielsen, K Sneppen, Self-inhibiting percolation and viral spreading in epithelial tissue. eLife 13 (2024).

26. S Peterl, et al., Morphology-dependent entry kinetics and spread of influenza a virus. The EMBO J. 44, 3959–3982 (2025).

27. EA Voigt, A Swick, J Yin, Rapid induction and persistence of paracrine-induced cellular antiviral states arrest viral infection spread in a549 cells. Virology 496, 59–66 (2016).

28. BP Holder, et al., Assessing the in vitro fitness of an oseltamivir-resistant seasonal a/h1n1 influenza strain using a mathematical model. PLoS ONE 6, e14767 (2011).

29. L Zhang, ME Peeples, RC Boucher, PL Collins, RJ Pickles, Respiratory syncytial virus infection of human airway epithelial cells is polarized, specific to ciliated cells, and without obvious cytopathology. J. Virol. 76, 5654–5666 (2002).

30. SN Roach, et al., Tropism for ciliated cells is the dominant driver of influenza viral burst size in the human airway. Proc. Natl. Acad. Sci. United States Am. 121, e2320303121 (2024).

31. NG Ravindra, et al., Single-cell longitudinal analysis of sars-cov-2 infection in human airway epithelium identifies target cells, alterations in gene expression, and cell state changes. PLOS Biol. 19, e3001143 (2021).

32. (2026) Created in BioRender. Hvid, U. https://BioRender.com/y8gfb2k.

33. Wikipedia contributors, Volterra integral equation (https://en.wikipedia.org/wiki/Volterra_integral_equation) (2026) Wikipedia, The Free Encyclopedia. Accessed: 2026-04-10.

34. T Bazett, Section 3.3: Convolution (https://web.uvic.ca/∼tbazett/diffyqs/convolution_section.html) (2026) Section 3.3 of an online textbook adapted from Jiří Lebl’s Notes on Diffy Qs. Accessed: 2026-04-10.

35. H Miller, The pole diagram and the Laplace transform (MIT 18.03 Supplementary Notes, Chapter 27) (n.d.) Accessed: April 15, 2026.

36. J Wallinga, M Lipsitch, How generation intervals shape the relationship between growth rates and reproductive numbers. Proc. Royal Soc. B: Biol. Sci. 274, 599–604 (2006).

37. AJ Lotka, Relation between birth rates and death rates. Science 26, 21–22 (1907).

38. Wikipedia contributors, Euler–Lotka equation (2026) Wikipedia, The Free Encyclopedia. Accessed: 2026-03-23.

39. DF Young, et al., Virus replication in engineered human cells that do not respond to interferons. J. Virol. 77, 2174–2181 (2003).

40. ME Becker, et al., Live imaging of airway epithelium reveals that mucociliary clearance modulates sars-cov-2 spread. Nat. Commun. 15 (2024).

41. S Chen, et al., Heterocellular induction of interferon by negative-sense rna viruses. Virology 407, 247–255 (2010).

42. CF Hatton, et al., Delayed induction of type i and iii interferons mediates nasal epithelial cell permissiveness to sars-cov-2. Nat. Commun. 12 (2021).

43. AB Russell, E Elshina, JR Kowalsky, AJW te Velthuis, JD Bloom, Single-cell virus sequencing of influenza infections that trigger innate immunity. J. Virol. 93 (2019).

44. EA Thayer, et al., Single-cell heterogeneity in interferon induction potential is heritable and governed by variation in cell state. bioRxiv (2025) Preprint.

45. KK Au, et al., Tracking the transcription kinetic of sars-cov-2 in human cells by reverse transcription-droplet digital pcr. Pathogens 10, 1274 (2021).

46. EW Weisstein, Lambert W-function (MathWorld —A Wolfram Resource) (n.d.) Last updated March 25, 2026. Accessed April 22, 2026.

47. N Khandelwal, et al., Studies on growth characteristics and cross-neutralization of wild-type and delta sars-cov-2 from hisar (india). Front. Cell. Infect. Microbiol. 11 (2021).

48. T Frensing, et al., Influenza virus intracellular replication dynamics, release kinetics, and particle morphology during propagation in mdck cells. Appl. Microbiol. Biotechnol. 100, 7181–7192 (2016).

49. M Diefenbacher, J Sun, CB Brooke, The parts are greater than the whole: the role of semi-infectious particles in influenza a virus biology. Curr. Opin. Virol. 33, 42–46 (2018) Virus vector interactions • Special Section: Multicomponent viral systems.

50. B Israelow, et al., Mouse model of sars-cov-2 reveals inflammatory role of type i interferon signaling. J. Exp. Medicine 217 (2020).

51. PP Ogger, et al., Type i interferon receptor signalling deficiency results in dysregulated innate immune responses to sars-cov-2 in mice. Eur. J. Immunol. 52, 1768–1775 (2022).

52. JM Rojas, A Alejo, V Martín, N Sevilla, Viral pathogen-induced mechanisms to antagonize mammalian interferon (ifn) signaling pathway. Cell. Mol. Life Sci. 78, 1423–1444 (2020).

53. E Makuch, I Jasyk, A Kula, T Lipiński, J Siednienko, Ifn-induced cxcl10 chemokine expression is regulated by pellino3 ligase in monocytes and macrophages. Int. J. Mol. Sci. 23, 14915 (2022).

54. KL Oslund, et al., Synergistic up-regulation of cxcl10 by virus and ifn in human airway epithelial cells. PLoS ONE 9, e100978 (2014).

55. U Rand, et al., Multi-layered stochasticity and paracrine signal propagation shape the type-i interferon response. Mol. Syst. Biol. 8 (2012).

56. V Schirrmacher, Important role of interferon regulatory factor (irf)-3 in the interferon response of mouse macrophages upon infection by newcastle disease virus. Int. J. Oncol. (2011).

57. DP Nayak, RA Balogun, H Yamada, ZH Zhou, S Barman, Influenza virus morphogenesis and budding. Virus research 143, 147–161 (2009).

58. P V’kovski, A Kratzel, S Steiner, H Stalder, V Thiel, Coronavirus biology and replication: implications for sars-cov-2. Nat. Rev. Microbiol. 19, 155–170 (2021).

59. A Timm, J Yin, Kinetics of virus production from single cells. Virology 424, 11–17 (2012).

60. OS Lund, U Hvid, BF Nielsen, K Sneppen, Stochasticity in viral infection and host response: A competition between speed and reliability. bioRxiv (2026) Preprint.

61. A Yakimovich, et al., Cell-free transmission of human adenovirus by passive mass transfer in cell culture simulated in a computer model. J. Virol. 86, 10123–10137 (2012).

62. MD Vahey, DA Fletcher, Influenza a virus surface proteins are organized to help penetrate host mucus. eLife 8, e43764 (2019).

63. JL McAuley, BP Gilbertson, S Trifkovic, LE Brown, JL McKimm-Breschkin, Influenza virus neuraminidase structure and functions. Front. Microbiol. 10 (2019).

64. SR Paludan, et al., Early host defense against virus infections. Cell Reports 43, 115070 (2024).

65. U Hvid, LEAP — Lotka-Euler Airway Pathogen model (https://github.com/ulrikhvid/ LEAP---Lotka-Euler-Airway-Pathogen-model) (2026) Code repository for Long-range seeding drives exponential growth in early respiratory viral infection.

