## Supplemental figures and derivations for "Bridging spatial scales in respiratory viral infection"

#### This PDF file includes:

Figs. S1 to S2
SI References

### 1. Supplementary material

**A. Area contribution.** The result that the mean area $\langle A \rangle = 1/q$ in the exponential growth phase turns out to be independent of the specific birth kernel, and follows solely from the relation between area and birth rate,

$$b(a) = q \cdot \frac{\mathrm{d}A}{\mathrm{d}a}, \tag{S1}$$

and the general Lotka–Euler equation,

$$\int_0^\infty b(a) e^{-\lambda a} \mathrm{d}a = q \int_0^\infty \mathrm{d}A \, e^{-\lambda a} = 1. \tag{S2}$$

Integrating by parts,

$$q \left[ A(a) \, e^{-\lambda a} \Big|_0^\infty + \lambda \int_0^\infty A(a) \, e^{-\lambda a} \, \mathrm{d}a \right] = 1. \tag{S3}$$

Since $A(0) = 0$ and $A(a)$ grows slower than exponentially, the first term vanishes. Since $P(a) = \lambda e^{-\lambda a}$, the remaining integral is simply $\langle A \rangle$, giving

$$q \cdot \langle A \rangle = 1 \implies \langle A \rangle = \frac{1}{q}. \tag{S4}$$

**B. Exact representation of stochastic IFN activation.** In the previous representation of the model, we made the simplifying assumption that IFN always gets activated at precisely the time when $A = A_n = 1/p_n$. In reality, activation is stochastic, and it is slightly more exact to treat it as a Poisson process. Then we can change $I \to I_2$, defined as

$$I_2(\lambda; a_g, a_n) = \int_{a_g}^\infty e^{-\lambda a} \cdot (a - a_g) \cdot e^{-A(a)/A_n} \mathrm{d}a = e^{-\lambda a_g} \int_0^\infty e^{-\lambda t} \cdot t \cdot e^{-\pi(v \; t)^2 p_n} \mathrm{d}t. \tag{S5}$$

with $t \equiv a - a_g$, $A(a) = \pi(v \; a)^2$ and $A_n = 1/p_n$. The solution is

$$I_2(\lambda; a_g, a_n) = e^{-\lambda a_g} \left[ \frac{1}{2\pi v^2 p_n} - \frac{\lambda}{4\pi v^3 p_n^{3/2}} \, e^{\frac{\lambda^2}{4\pi v^2 p_n}} \, \mathrm{erfc}\left( \frac{\lambda}{2\sqrt{\pi} \, v \sqrt{p_n}} \right) \right] \tag{S6}$$

### References


1. N Khandelwal, et al., Studies on growth characteristics and cross-neutralization of wild-type and delta sars-cov-2 from hisar (india). *Front. Cell. Infect. Microbiol.* **11** (2021).

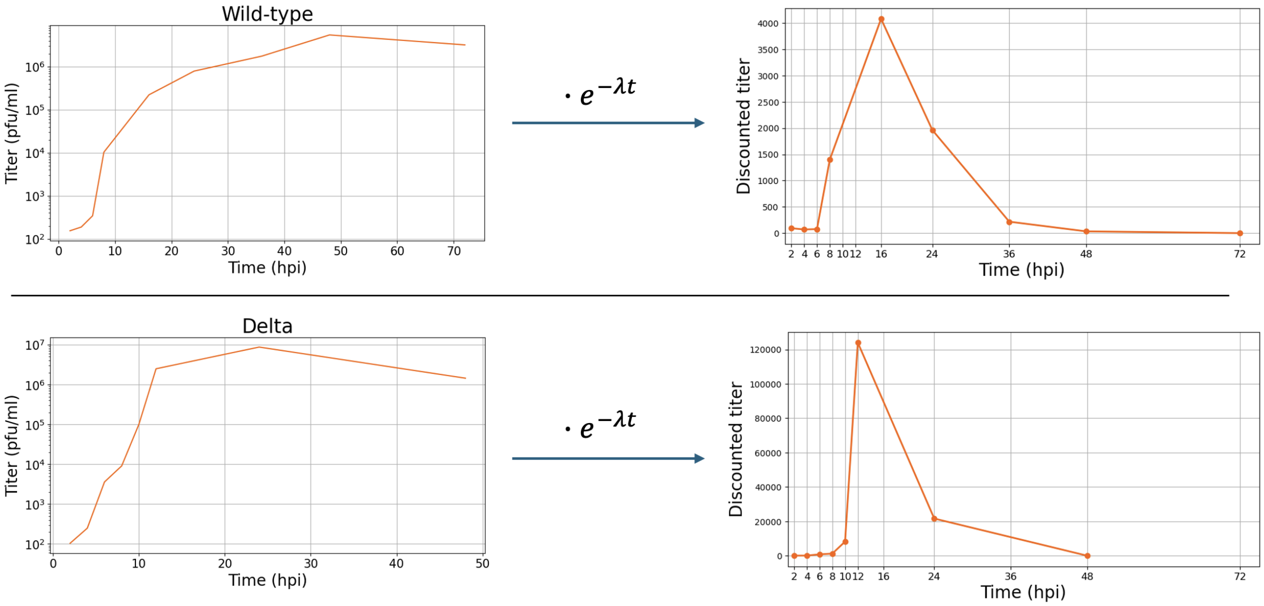


**Fig. S1. Applying exponential discount to one-step growth curves yields $a_g$ estimate** Left: Replotting of one-step growth curves from (1). These curves are titer measurements from cultures infected at high MOI, such that they represent the output of just one viral generation. Data collected from figures in (1) with graphreader.com. Right: Discounting late releases by multiplying by an exponential function with $\lambda = 0.25\ \mathrm{h}^{-1}$ yields shapes with clear peaks at 16 and 12 hpi respectively. These are our estimates of $a_g$ for SARS-CoV-2 WT and Delta.

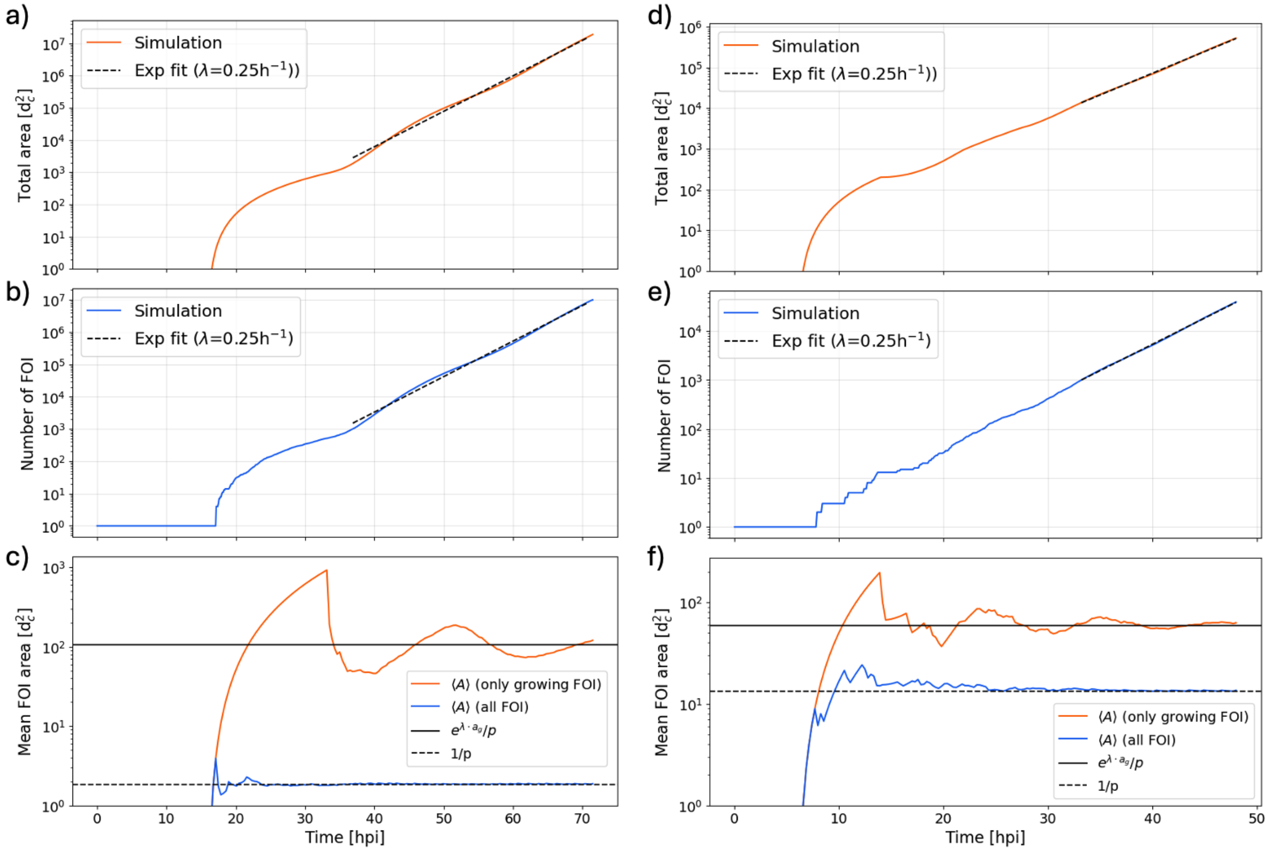


**Fig. S2. Stochastic simulation of the birth kernel shows transient and exponential phase.** Simulation using the parameter values in Table 1 with **a), b), c)** using SARS-CoV-2 parameters and **d), e), f)** using parameters for IAV. Both total area **a)** and number of IF **b)** show a transient period with oscillations on the timescale of $a_g$ h, before settling into steady exponential growth with $\lambda = 0.25$/h as predicted analytically. Panel **c)** shows the mean value of all IF (blue) and all growing IF (red) respectively. As non-growing IF will be invisible or tiny in a measurement using fluorescent proteins, the red curve is more representative of what might actually be observed, i.e., a typical focus size of $\approx 100$ cells.

 **Ulrik Hvid, Bjarke Frost Nielsen, Kim Sneppen**
